# The role of the VirB ligand CTP in the molecular mechanism of transcriptional anti-silencing in *Shigella flexneri*

**DOI:** 10.64898/2025.12.19.695565

**Authors:** Taylor M. Gerson, Monika MA. Karney, Helen J. Wing

## Abstract

In bacteria, nucleoid-structuring proteins bind and constrain DNA, often leading to transcriptional silencing. In *Shigella spp*., the histone-like nucleoid-structuring protein H-NS silences many genes on the large virulence plasmid. Upon a shift to human body temperature, VirB, a DNA-binding protein and key transcriptional regulator of the *Shigella* virulence cascade, is produced. VirB counteracts H-NS-mediated transcriptional silencing and belongs to a fast-evolving clade of the ParB superfamily. Like other ParB proteins, VirB binds the ligand CTP. While CTP is essential for the anti-silencing activity of VirB, the role of CTP in the mechanism of VirB-dependent anti-silencing has yet to be determined. This work shows that VirB does not require CTP for specific engagement of its DNA recognition site, but CTP is necessary for the formation of large VirB-DNA complexes, which likely form when VirB dimers adopt a sliding clamp conformation. Furthermore, CTP-binding mutants do not trigger a loss of negative supercoils from plasmid DNA, a VirB-dependent activity proposed to destabilize H-NS-DNA complexes. Together, our findings provide novel insight into the relationship between VirB, CTP, and DNA and reveal the role of the VirB ligand, CTP, in regulating *Shigella* virulence.

## Introduction

In bacteria, nucleoid-structuring proteins (NSPs) such as Fis, IHF, H-NS, HU, and LRP, impact gene expression (1). NSPs bind DNA to remodel, organize, and constrain the genetic material inside bacterial cells. In some cases, these NSP-DNA interactions transcriptionally silence parts of the genome (1–4). Silenced genes can be upregulated through a process called anti-silencing, where the NSPs responsible for silencing gene expression are offset. These opposing processes occur in response to new environmental stimuli and influence bacterial adaptation. Anti-silencing, therefore, regulates the transcription of stress response genes (5) and key virulence genes in many important bacterial pathogens (6–8). Mechanistically, anti-silencing can occur through the use of DNA-binding proteins called anti-silencers. A diverse range of anti-silencers has been identified (VirB (9), SlyA (10), RovA (11), ToxT (12), RegA (13), LeuO (14,15), etc.), which has led to the hypothesis that several solutions may exist to help offset NSP-mediated silencing of genes (16). Despite the lack of homology between these anti-silencing proteins, however, it is conceivable that the underlying molecular mechanism of transcriptional anti-silencing is conserved.

In *Shigella* species, the causative agents of bacillary dysentery (17–20), at 30°C, the histone-like nucleoid structuring protein, H-NS, silences many virulence genes encoded on the large virulence plasmid (21–23). Subsequently, upon a temperature shift to 37°C, the virulence cascade is activated, leading to the synthesis of a key transcriptional regulator of *Shigella* virulence, VirB (24–26). VirB is a bona fide anti-silencing protein that counteracts H-NS-mediated silencing of *Shigella* virulence genes (27). This is supported by the following: i) in the absence of H-NS, VirB has little to no effect on gene expression (26); ii) *Shigella* strains lacking *virB* are completely avirulent, because a shift to 37°C alone is not sufficient to relieve H-NS-mediated silencing (26); and iii) if VirB is artificially produced at 30°C, VirB still alleviates H-NS-mediated silencing, even at this non-permissive temperature.

Interestingly, VirB is not evolutionarily related to other transcriptional regulators. Instead, VirB is a member of the ParB superfamily (9,28,29). Traditionally, ParB proteins are characterized to contribute to the segregation of chromosomal and plasmid DNA to the cell pole before cell division (30–32). To function in this process, canonical ParB members (*Bacillus subtilis* Spo0J (33), *Bacillus subtilis* Noc (34), *Myxococcus xanthus* ParB (35), *Myxococcus xanthus* PadC (35), *Caulobacter vibrioides* ParB (36), *Corynebacterium glutamicum* ParB (37), *Streptomyces coelicolor* ParB (38), etc.) bind cytidine triphosphate (CTP) as a dimer, and recognize a DNA recognition site, known as the *parS site*. Dimeric ParBs bound to CTP and *parS* adopt a DNA sliding clamp conformation, so the DNA double helix drops into the central lumen of the dimer, allowing ParB to dissociate from *parS.* The sequential recruitment of additional ParB dimers to the now-liberated *parS* site triggers the bidirectional spread of ParB along DNA, like beads on a string (33,35,36).

While VirB is a member of the ParB superfamily, VirB does not function in DNA segregation but rather acts as a key transcriptional regulator of *Shigella* virulence. Transcriptional anti-silencing by VirB relies on three key steps: i) the specific binding of dimeric VirB to its DNA recognition site (39); ii) the spreading of VirB dimers along the DNA (40); and iii) the VirB-mediated loss of negative supercoils in the region bound by H-NS (41). The need for VirB to bind its DNA recognition site to accomplish its regulatory activities comes from site-directed mutagenesis of the VirB binding site and DNA-binding mutants, which are unable to anti-silence (9,25,39). The spreading of VirB dimers on DNA is supported by the ParB literature and DNA roadblocking studies at *PicsP* (40), where a DNA roadblock positioned on the downstream but not upstream flank of the VirB binding site prevents transcriptional anti-silencing (40). Thus, the unidirectional spread of VirB along the DNA helix towards the region occupied by H-NS appears necessary for VirB-dependent transcriptional anti-silencing. Finally, following engagement of its binding site *in vivo* and *in vitro,* VirB causes a loss of negative supercoils from plasmid DNA (41). These topological changes are transient in nature, but have been implicated in the mechanism of transcriptional anti-silencing because a loss of negative supercoils in the vicinity of a region bound by H-NS can alleviate H-NS-mediated silencing, even in the absence of VirB (41).

Our recent work (29), as well as that of others (42,43), reveals that, like chromosomal ParB proteins, VirB binds CTP preferentially and with specificity over other NTP ligands, such as ATP, GTP, and UTP. Furthermore, although UTP is the NTP most closely related to CTP, UTP does not bind to VirB, as UTP does not generate a VirB binding signature in isothermal titration calorimetry experiments (29). VirB mutants unable to bind CTP, including VirB T68A and R94A are incapable of transcriptional anti-silencing, whereas mutants with substitutions in these or neighboring residues (T68S or R95A) that retain CTP binding retain wild-type anti-silencing activity (29). Thus, CTP is necessary for the VirB-dependent regulation of virulence genes (29). While steps required for VirB-dependent anti-silencing have been delineated, it remains unclear what role CTP plays in each step. Therefore, the overarching goal of this study is to determine how CTP governs VirB-DNA interactions by examining which of the three steps of VirB-dependent anti-silencing, DNA binding, spreading, and modulation of DNA supercoiling require CTP.

## Material and Methods

### Bacterial strains, plasmids, and media

The strains and plasmids used in this study are listed in Table S1 *E. coli* strains were routinely grown on LB agar (LB broth containing 1.5% [wt vol^-1^] agar). Liquid cultures were grown overnight at 37°C in LB broth. Overnight cultures were diluted 1:100 and subcultured at 37°C with aeration in the specified media. To ensure plasmid maintenance, antibiotics were used at the following final concentrations: ampicillin, 100 µg mL^−1^, and chloramphenicol, 25 µg mL^−1^.

### VirB Protein Purification

Wild-type VirB was purified by the Monserate Biotechnology Group (San Diego, CA) as previously described in (41).

### Linear DNA Electromobility Shift Assay

Electromobility shift assays were used to test VirB-DNA binding *in vitro*. For linear DNA fragments, EMSA was performed as described in (40). Briefly, two 54 bp *icsP* promoter fragments containing either a wild-type or mutated VirB binding site were created by annealing primer pairs W391/W392 and W393/W394, respectively. Primer pairs were denatured at 95°C for 5 min and further annealed using a cycle that decreases by 1°C every minute until reaching a final temperature of 5°C. Each annealed DNA product was gel-purified on 6% non-denaturing PAGE and electroeluted, and the DNA was extracted by phenol-chloroform and ethanol precipitated. DNA was single-end radiolabeled using T4 polynucleotide kinase (Promega Cat. No. M4101) using [γ^32^P] ATP (specific activity, 3000 Ci mmol^−1^; Perkin Elmer). Unincorporated radionucleotides were removed by G-50 Micro Columns according to the manufacturer’s directions (GE Healthcare). Varying concentrations of purified VirB-His_6_ (Monserate Biotechnology Group) in a 1X VirB protein buffer (25 mM Hepes pH 7.6, 100 mM NaCl, 0.1 mM EDTA, 0.5 mM BME, 5% glycerol) were incubated with 5 nM of each *icsP* promoter target in 1X EMSA binding buffer (50 mM Tris-HCl pH 8.0, 20 mM KCl, 10% glycerol, 1000 μg μL^-1^ BSA, 400 μg μL^-1^ Herring Sperm DNA), in a total volume of 20 μL, at 37°C for 20 min. Reactions were resolved on a 6% non-denaturing PAGE in 1X TBE at 60 V for 3 hours. The gel was transferred to Whatmann paper, covered in plastic wrap, and exposed to a storage phosphor screen (Kodak). The screen was imaged on an Amersham Typhoon.

### Quantification of Linear Electromobility Shift Assays

Free DNA bands were quantified using the ‘Analysis Toolbox’ in the AzureSpot Analysis Software version 2.0.062. Lanes were identified using the ‘create lanes’ feature, and boxes were drawn manually with automatic edge detection using the ‘detect bands’ feature. Band density (in pixels) of the free DNA band was determined following background subtraction using the rolling ball method (radius 200), and the resulting stacked lane profiles are presented. The resulting pixel values were used to normalize the data relative to the no-protein lane in each test condition and were reported as percentages. The normalized percentage of the free DNA band (bottom of the gel) was graphed and is shown in Fig. 1B and S1B.

**Figure 1.**
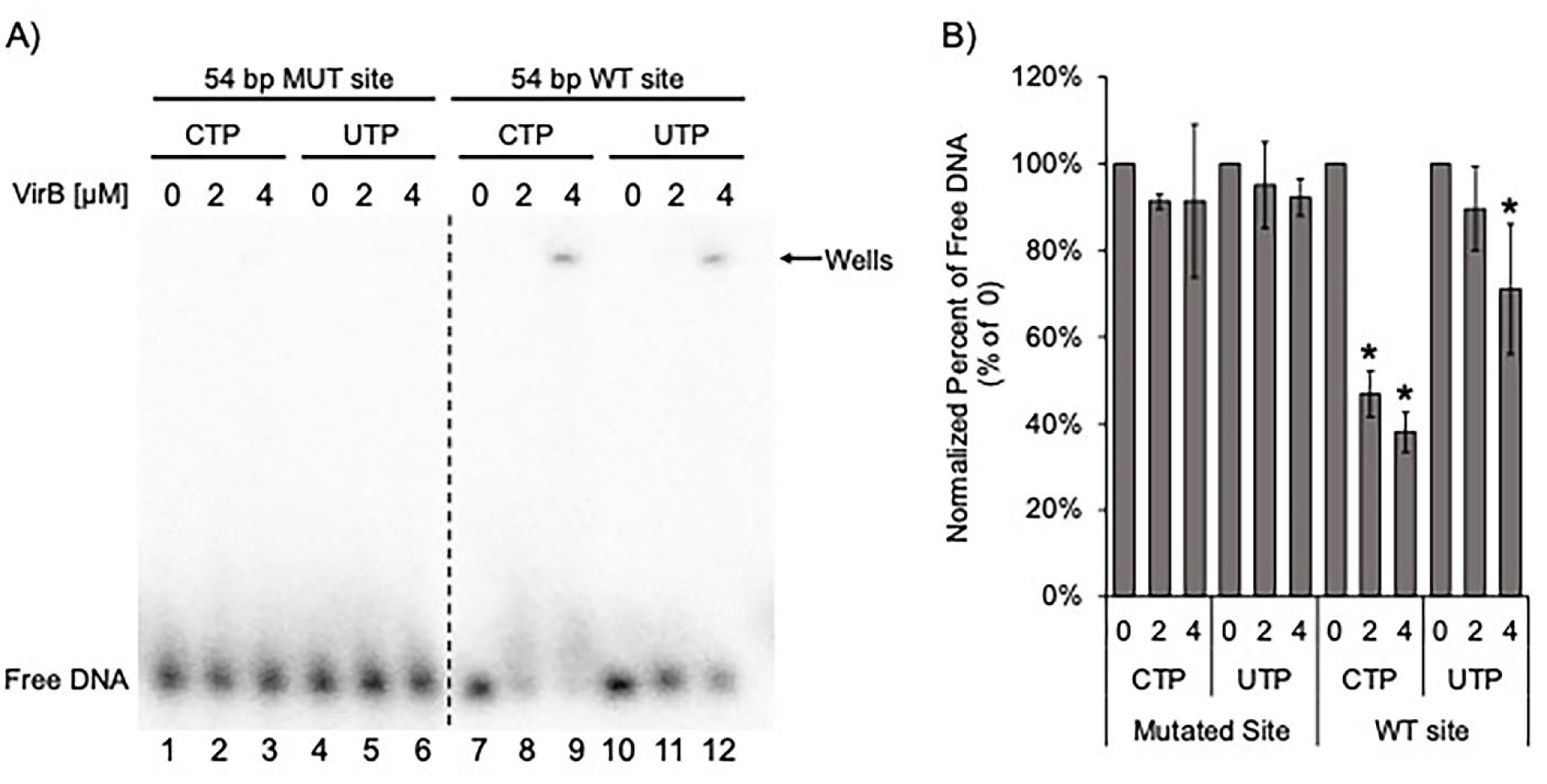
*In vitro,* CTP is not required for specific DNA binding by VirB. A) Electromobility shift assay using 54 bp linear DNA containing either a WT or mutated VirB binding site, incubated with either CTP or the negative control UTP (VirB does not bind UTP, determined by ITC (29)), and increasing concentrations of VirB (representative image shown). B) Quantification of signal loss of the lower band as VirB concentration increases (n=3). Significance was calculated using a one-way ANOVA post-hoc Tukey HSD, p<0.05. Asterisks, *, indicate a significant decrease compared to 0 µM lanes in each given condition. Complete statistical analysis provided in Table S2. **ALT TEXT:** Representative EMSA gel showing DNA shifts and bar graph showing quantification (n=3). Visible shifts appear in lanes containing wild-type VirB binding site in the presence of either CTP or UTP, while lanes with the mutated VirB binding site show no visible DNA shift. These data indicate VirB does not require CTP for its specific engagement of the VirB recognition site.

### Plasmid DNA Electromobility Shift Assay

To evaluate the importance of CTP in VirB binding and formation of large VirB-DNA complex on plasmid DNA, a modified electrophoretic mobility shift assay (41,44) using the P*icsP-lacZ* plasmid (pHJW20; (26)) was performed. The plasmid pHJW20 was prepared by isolating from DH10B cells using the Promega Pure Yield Plasmid Miniprep System (Promega #A1223). To prepare the DNA stock, 3 mL of culture was processed according to the manufacturer’s instructions using a mini-prep method, and the resulting mini-preps were combined. A fixed volume of the DNA stock (30 μL) was incubated with increasing concentrations of VirB (0.1175 μM, 0.235 μM, 1.175 μM) for 30 min at 37 °C with either CTP or UTP (1mM; NEB #N0450S). Each reaction contained 1x VirB buffer (25 mM HEPES, pH 7.6, 100 mM NaCl, 0.1 mM EDTA, 0.5 mM β-mercaptoethanol, 5% glycerol, 5 mM MgCl_2_). After incubation with VirB, 5 μL was removed from each sample to analyze VirB binding to plasmid DNA using electrophoresis (0.7% agarose gel in 1× TAE, 100 V for 1 h, and imaged after staining with ethidium bromide). High molecular weight complexes that form in/near the well may represent non-specific interactions due to the lack of competitor DNA and/or carrier proteins in these reactions.

### VirB-DNA capture assay

After verifying VirB binding to the DNA via plasmid EMSA, samples were incubated at 65°C for 30 min to denature VirB. Phenol/chloroform and ethanol precipitation were used to isolate DNA not bound by VirB. Free DNA and VirB will separate into the aqueous and organic layers, respectively. Therefore, any DNA still bound or “captured” by VirB would also be trapped in the organic layer and lost during recovery. The recovered free DNA is visualized after gel electrophoresis on agarose and staining with ethidium bromide.

### *In vivo* supercoiling assay

For *in vivo* supercoiling assays, plasmid DNA was extracted from *E.coli* DH10B cells harboring the P*icsP*-*lacZ* reporter and the pBAD-*virB* expression plasmid or its derivatives as described in (41,44). Briefly, DH10B carrying double plasmid or single plasmid controls was grown overnight in LB supplemented with the appropriate antibiotics and with 0.2% (wt. vol^−1^) D-glucose (45) if the strain contained pBAD-*virB* or its derivatives. Following overnight growth, cells were subcultured by diluting 1:100 in LB broth with the appropriate antibiotics and grown for 3 h at 37°C. VirB expression was induced in half of the cultures with 0.2% (wt. vol^−1^) L-arabinose, and all cultures were grown for an additional 2 h at 37°C. Following growth, cells were harvested, normalized to cell density (OD_600_ nm), and DNA was isolated using the PureYield Plasmid Miniprep System (Promega, Cat. No. A1222) according to the manufacturer’s instructions.

The topoisomer distribution of the isolated DNA samples was analyzed using 1D chloroquine gel electrophoresis as described in (44). Routinely, 500 ng of DNA was electrophoresed for samples containing both plasmids, and 250 ng was electrophoresed for single-plasmid controls. Samples were electrophoresed through a 2× TBE [178 mM Tris base, 178 mM borate, 4 mM EDTA (pH 8.0)], 0.7% (wt. vol^−1^) agarose gel, for 24-26 h in the dark at 2.0–2.5 V cm^−1^ in the presence of chloroquine at a final concentration of 2.5 μg mL^−1^. After electrophoresis, gels containing chloroquine were destained in sterilized distilled water, changing the water every 30–60 min. After destaining, gels were stained with ethidium bromide, imaged using a UVP BioSpectrum 410 Imaging System, and the resulting images were manipulated using Visionworks™ LS Image Acquisition.

### Quantification of Plasmid EMSA, VirB-DNA Capture Assay, and *in vivo* Supercoiling Assay

Topoisomer bands were quantified using the ‘Analysis Toolbox’ in the AzureSpot Analysis Software version 2.0.062 as described in (41). Briefly, an initial line was drawn upwards through the center of the lanes using the ‘draw shapes’ feature. Subsequent lines were copied using the ‘duplicate’ feature and dragged to adjacent lanes. The resulting gel images and lane profiles, which indicate the density of each topoisomer band, were exported and are shown in Figs. 2B, 2D, S2, S4.

**Figure 2.**
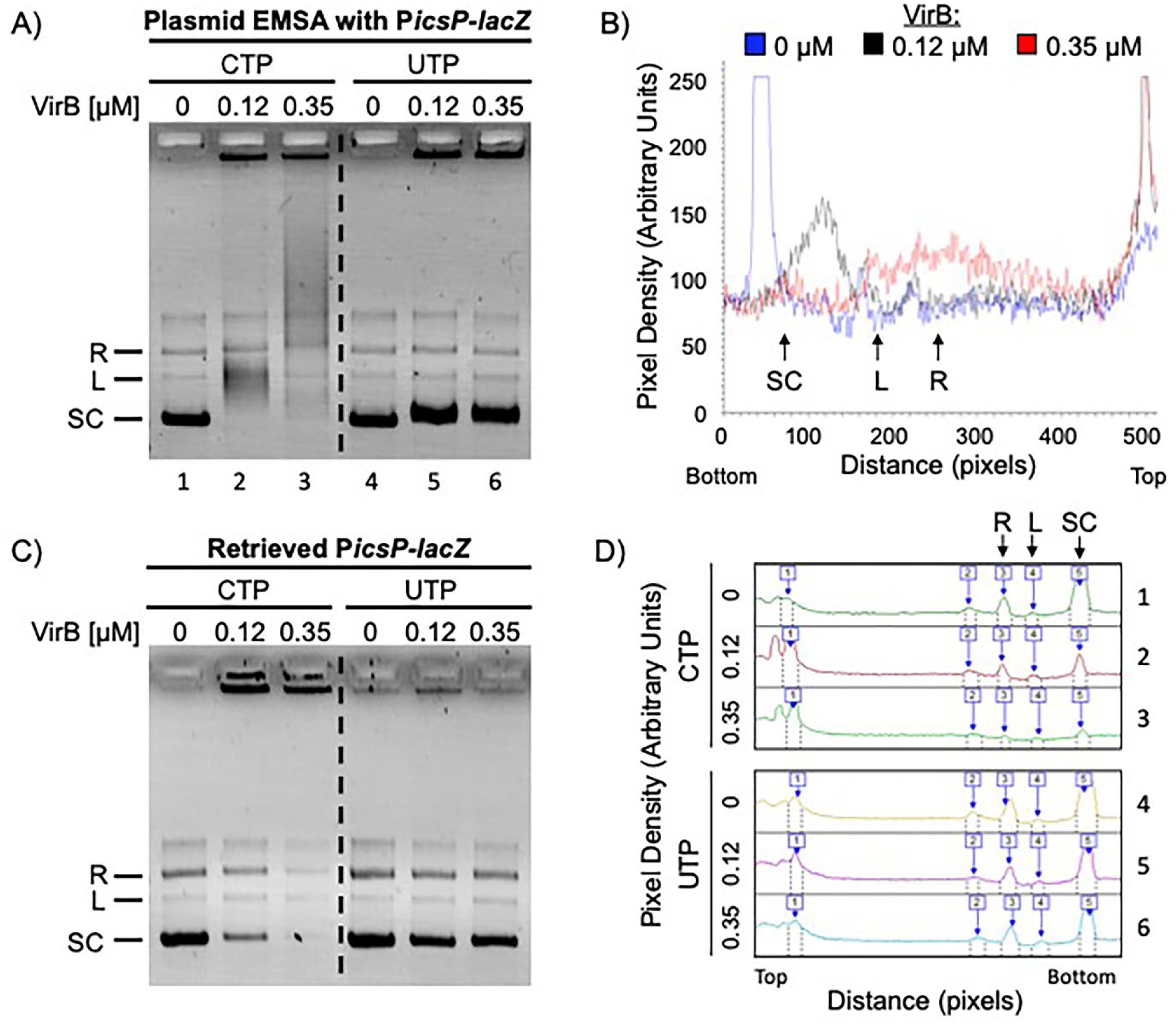
*In vitro,* CTP is necessary for the formation of large VirB-DNA complexes that are resistant to heat dissociation. A) Modified electromobility shift assay with plasmid DNA containing a VirB binding site (P*icsP-lacZ*) with excess CTP or UTP (1 mM) and increasing concentrations of VirB (0, 0.12, 0.35 μM; representative image shown). B) Lane profile of the shift of P*icsP-lacZ* in the presence of CTP (lanes 1-3) from Fig. 2A is shown (lane profile for UTP provided in Fig. S2). The molar ratio of VirB to DNA required to shift plasmid DNA was calculated to be 29:1. C) DNA retrieved from plasmid EMSAs after heat treatment, phenol/chloroform extraction, and ethanol precipitation (representative image shown). D) Lane profiles of DNA retrieved from reactions shown in panel C, incubated with either CTP (top) or UTP (bottom). Peaks corresponding to supercoiled (SC), linear (L), and relaxed (R) bands are indicated. **ALT TEXT:** Gels and quantification line graphs, labelled A to D. A, EMSA gel with plasmid DNA showing DNA shifts in the presence of CTP or UTP. B, lane trace of CTP-containing samples showing an incremental shift in the supercoiled DNA population as VirB concentrations increase. C, agarose gel of DNA recovered from plasmid EMSA after heat treatment, phenol/chloroform extraction, and ethanol precipitation, where a visible loss of DNA occurs in samples containing CTP, but not UTP. D, lane trace of panel C, showing striking loss of supercoiled DNA in samples with CTP, but not UTP.

### Calculating the VirB:DNA Ratio in Electromobility Shift Assays

To determine the ratio of VirB to the DNA substrate required to observe a shift in linear and plasmid electromobility shift assays, the molar concentrations of both VirB and DNA were calculated. For DNA, the total nanograms used in the assay were converted to molarity based on the length of the dsDNA, assuming 1 base pair is equivalent to 660 Da. For VirB, the final concentration in each reaction was calculated from the purified stock’s starting concentration (94 mM). Molar concentration of VirB and the DNA were compared to one another within a given assay, and the resulting ratios are listed in Fig. S1B and 2.

### Statistical Analyses

All statistical calculations were performed using IBM SPSS Statistics for Windows (version 28.01.0) as described in (29). Routinely, One-way ANOVA tests were used, and *post hoc* analyses were performed as indicated in the figure legends. Throughout this work, statistical significance is denoted *, indicating p<0.05.

## Results

### *In vitro,* VirB does not require CTP for engagement of its DNA-binding site

To determine the role that CTP plays in VirB-DNA interactions, we first determined whether CTP was necessary for VirB to specifically bind to a DNA fragment bearing its DNA-binding site *in vitro*. Using an electromobility shift assay (EMSA), reaction mixes containing purified VirB protein were incubated with a radiolabeled 54 bp linear DNA fragment containing either a wild-type or mutated VirB binding site in the presence of CTP or UTP (UTP serves as a negative control, as it does not bind to VirB (29)). As expected, regardless of whether CTP or UTP was present, when increasing concentrations of VirB were incubated with DNA fragments containing a mutated VirB binding site, no dramatic shifts in the DNA were observed (Fig. 1A-B; lanes 1-6). In contrast, when increasing concentrations of VirB were incubated with DNA fragments bearing the wild-type VirB binding site, shifts of the target were observed in the presence of both CTP and UTP (Fig. 1A-B; lanes 7-12). Even though our reaction mixes included competitor carrier protein and non-specific DNA, the shifts did not appear as discrete protein-DNA complexes (the top band represents complexes that remain in the well) but rather appeared as a smear up the gel, likely representing heterogeneous VirB-DNA complexes. Consequently, to quantify these data, we measured the loss of the free DNA target signal as VirB concentrations increased.

In the presence of a wild-type VirB binding site and in the presence of CTP, a significant loss in the amount of free DNA was observed as the concentration of VirB increased (Fig. 1A, lanes 7-9 and Fig. 1B). Similarly, in the presence of UTP, a loss of free DNA was observed, but even with the highest concentration of VirB used, the loss was not as pronounced as with CTP (Fig. 1A-B; lanes 10-12). Thus, VirB can bind to its specific target site *in vitro* in the absence of CTP, but its affinity for its DNA recognition site appears to be enhanced by CTP. These findings are further supported by additional experiments, where VirB (between 0-1 μM) was more finely titrated into reaction mixes (Fig. S1 and Table S3).

### *In vitro,* CTP is necessary for the formation of large VirB-DNA complexes

In *Shigella,* VirB binding sites are located on the large virulence plasmid (46,47), and evidence supports that VirB spreads along DNA after engaging its DNA recognition site (40,48). VirB spreading would lead to the formation of large VirB-DNA complexes on this closed circular DNA. Therefore, to determine the role of CTP in the formation of large VirB-DNA complexes, we next ran a modified EMSA using plasmid DNA (44). Purified VirB was incubated with a plasmid carrying the full-length *icsP* promoter, including the native VirB binding site, in either the presence of CTP or UTP. Since the plasmid DNA was not radiolabeled and DNA protein complexes were resolved on agarose gels, non-specific DNA could not be included in each reaction mix, and carrier protein was not included either. These modifications to standard EMSA reaction conditions may explain, to some extent, the large protein-DNA complexes observed near the wells in the presence of VirB (Fig. 2A, lanes 2-3 and 5-6). Regardless, in the presence of VirB and CTP, a dramatic shift in the DNA was observed (Fig. 2A, lanes 1-3). This shift appears as a smear that moves up the gel in a VirB concentration-dependent manner (Fig. 2B). In contrast, in the presence of UTP, a modest and discrete shift was observed (Fig. 2A), which remained unchanged with increasing VirB concentrations (Fig. S2). These data are consistent with the results of linear DNA EMSAs (Fig. 1), where VirB specifically engages its DNA recognition site in the absence of CTP (i.e., in the presence of the negative control, UTP). In contrast to the linear EMSAs, however, these data demonstrate that CTP is necessary for the formation of large VirB-DNA complexes on closed circular plasmid DNA. Additionally, these EMSAs also reveal that VirB preferentially binds to supercoiled plasmid DNA, because regardless of whether CTP or UTP is present, it is the supercoiled DNA band that shifts in the presence of VirB; the linear (L) and relaxed (R) DNA bands remain unchanged (Fig. 2A & B). This activity of VirB aligns well with its established role as a regulator of virulence plasmid genes.

To further characterize the nature of the large VirB-DNA complexes that form in our plasmid EMSAs, we next chose to evaluate the stability of the large complexes. While heat exposure typically dissociates DNA from DNA-binding proteins, we reasoned that DNA would be trapped by VirB complexes in the sliding clamp conformation. Such trapped plasmid:VirB complexes would partition with the protein fractions in phenol:chloroform extractions. Consequently, we incubated identical samples as those used in the plasmid EMSAs (Fig. 2A) at 65 °C for 30 minutes. The DNA was then extracted using phenol:chloroform and ethanol precipitation, and examined using agarose gel electrophoresis.

In the presence of CTP, the amount of plasmid DNA recovered was less as the concentration of VirB increased (Fig. 2C, lane 1-3, bands labeled R, L, & SC); this loss was most striking in the supercoiled band (Fig. 2D). In contrast, in the presence UTP, only a slight decrease in the amount of recovered plasmid DNA was observed (Fig. 2C, lanes 4-6), and this was the most striking with the highest concentration of VirB (Fig. 2D). Importantly, if VirB-DNA complexes were incubated with proteinase K before the phenol:chloroform extraction, equivalent amounts of DNA were retrieved, regardless of which NTP was present (data not shown). The inability to recover DNA from complexes formed in the presence of CTP is consistent with the large VirB-DNA complexes being locked on the DNA in a sliding-clamp conformation that is resistant to dissociation by heat. Intriguingly, the high molecular weight complexes running at the top of the gel, close to the wells in Fig. 2A (lanes 2-3 and 5-6), exhibited a recovery pattern opposite to the plasmid DNA: when formed with CTP, the high molecular weight complexes were retained, whereas when formed with UTP, they were not recovered. The reason for this remains unclear, but these findings demonstrate a clear difference between the high molecular weight complexes that form in the presence of CTP and UTP. In sum, the plasmid EMSAs and DNA retrieval assays demonstrate that the CTP ligand is required for the formation of large VirB-DNA complexes on plasmid DNA (seen in EMSAs (Fig. 2A)), and support that under these conditions, the VirB proteins are locked onto the plasmid DNA in sliding clamp conformation.

### VirB-dependent changes in DNA supercoiling *in vivo* are dependent on CTP binding

Our recent work has demonstrated that when VirB engages its DNA-binding site on plasmid DNA *in vivo,* a loss of negative supercoils is observed in the plasmid DNA (41). This VirB-dependent loss of negative supercoils has since been implicated in the mechanism of anti-silencing, because when a loss of negative supercoils is generated, independently of VirB, in a region bound by H-NS, it is sufficient to relieve the H-NS-mediated silencing of a nearby promoter (41). Thus, we next wondered whether CTP binding by VirB is necessary for VirB-mediated changes in DNA supercoiling.

To test this, we assessed whether VirB mutants unable to bind CTP caused a loss of negative supercoils from plasmid DNA bearing a VirB-binding site, using a previously described *in vivo* DNA supercoiling assay (41,44). To do this, we used L-arabinose inducible expression of *virB* alleles encoding VirB T68A or R94A, which are unable to bind CTP (Fig. S3A), but still capable of binding the VirB DNA recognition site (Fig. S3B), and VirB T68S or R95A, which retain CTP binding (Fig. S3A). Briefly, *E. coli* carrying P*icsP-lacZ* (which includes the native VirB binding site) were grown in the presence or absence of inducer. Following induction, plasmid DNA samples were isolated and electrophoresed in agarose gels containing chloroquine (2.5 μg mL^-1^), an intercalating agent that unwinds DNA by inserting between the strands of the double helix. It is well established that electrophoresis of plasmid DNA in the presence of chloroquine can better discern the topoisomeric state of the DNA (44).

In this experiment, all single plasmid controls (the pBAD-*virB* derivative (lane 1 & 14) and the plasmid carrying the VirB binding site (P*icsP-lacZ;* lanes 2 & 13)) showed little to no change in their topoisomer distribution regardless of the induction conditions (Fig. 3). As expected, in DNA samples containing both plasmids, the P*icsP-lacZ* reporter isolated from cells expressing a wild-type VirB displayed an altered topoisomer distribution near the top of the gel compared to P*icsP-lacZ* reporter isolated from cells without *virB* expression (Fig. 3; compare lanes 3 & 4). This was consistent with the previously reported VirB-dependent loss of negative supercoils from plasmid DNA (41). When P*icsP-lacZ* was isolated from cells expressing either VirB R95A or T68S, an altered topoisomer distribution matching that generated by wild-type VirB was observed (Fig. 3; compare lanes 7 & 8 and lanes 11 & 12 to lanes 3 & 4; quantification in Fig. S4 C & D). This was not surprising as these two VirB derivatives are both capable of binding CTP via DRaCALA (Fig. S3A). In contrast, the induction of either VirB R94A or T68A (two VirB derivatives unable to bind CTP via DRaCALA (Fig. S3A)) resulted in no change in the DNA topology of P*icsP-lacZ* (Fig. 3; compare lanes 5 & 6 and lanes 9 & 10 to lanes 3 & 4; quantification in Fig. S4 A & B). Thus, the VirB-dependent loss of negative supercoiling from plasmid DNA is entirely dependent on the ability of VirB to bind CTP.

**Figure 3.**
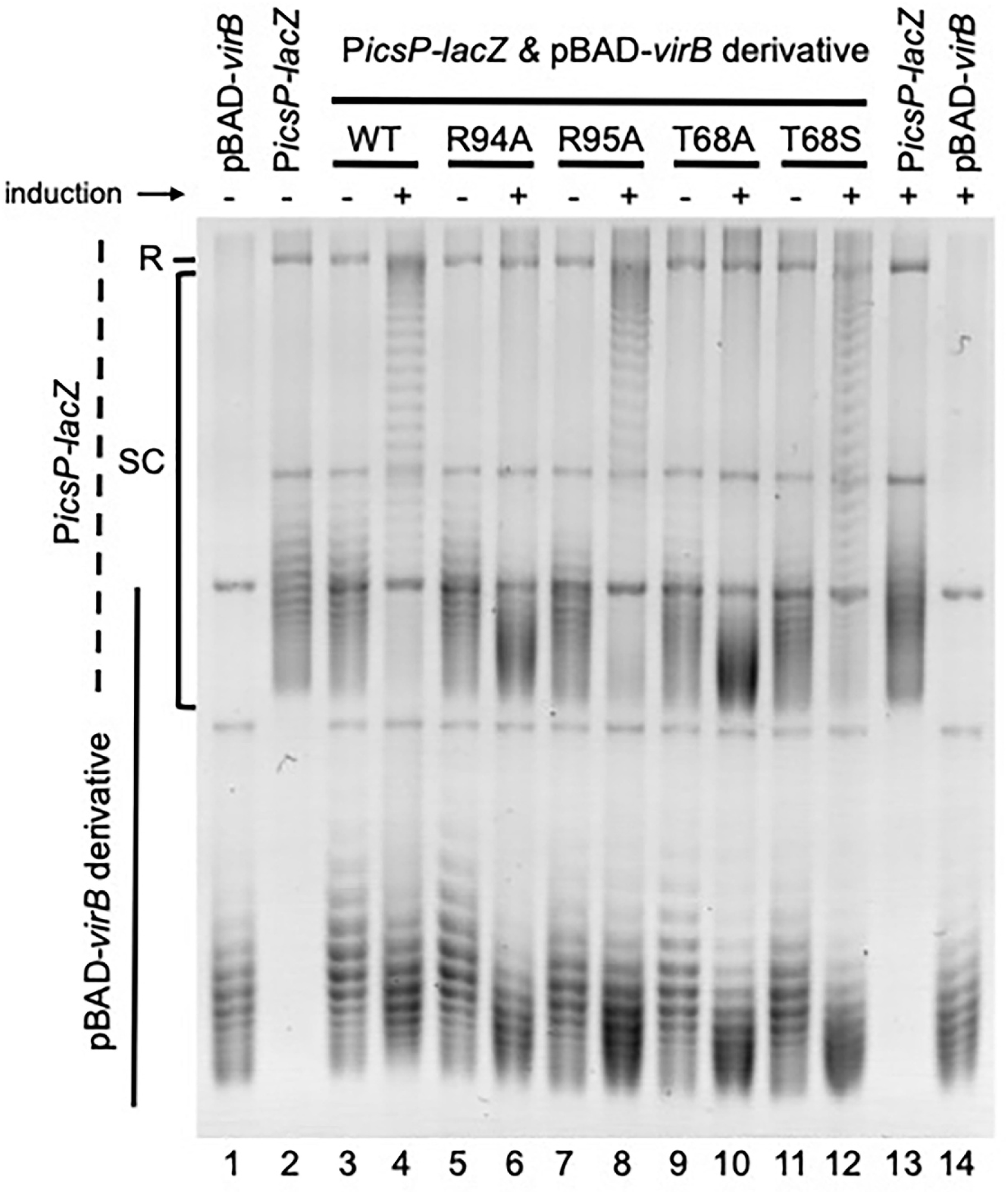
*In vivo* VirB-dependent changes in DNA supercoiling are dependent on CTP binding. Analysis of topoisomer distribution of P*icsP-lacZ* reporter isolated from *E. coli* in the presence or absence of wild-type or *virB* derivatives induced during growth on an agarose gel containing chloroquine (2.5 µg mL^-1^). Representative image shown. Quantification of key lanes (4, 6 & 10, and 4, 8 & 12) is provided in Fig. S4. **ALT TEXT:** Agarose gel showing changes in plasmid DNA topoisomer distributions. Altered distributions appear in lanes containing VirB derivatives capable of binding CTP, while lanes with VirB derivatives unable to bind CTP show unchanged topoisomer distributions. This demonstrates that CTP binding is required for VirB-dependent changes in DNA supercoiling.

## Discussion

In this work, we illuminate the role that CTP, the recently described ligand of VirB, plays in the mechanism of VirB-dependent anti-silencing. *In vitro*, CTP is not required for VirB binding of its DNA recognition site (Fig. 1), but is essential for the formation of large VirB-DNA complexes on plasmid DNA (Fig. 2A & B). Using a combination of plasmid EMSAs and DNA retrieval assays, the large VirB-DNA complexes that form on the DNA in the presence of CTP were found to be recalcitrant to heat dissociation (Fig. 2C & D). CTP binding by VirB is also required for the VirB-dependent modulation of DNA supercoiling (Fig. 3), an activity implicated in the relief of H-NS-mediated silencing. Consequently, our work establishes a connection between large VirB-DNA complex formation, VirB-dependent modulation of DNA supercoiling, and CTP for the first time, providing new insights into the mechanism of VirB-dependent transcriptional anti-silencing, a crucial regulatory process that controls *Shigella* virulence.

Our initial studies examined VirB-CTP-DNA interactions using EMSAs with traditional short linear DNA targets. VirB was found to bind its DNA recognition site, regardless of whether CTP was present or not (Fig. 1), although CTP did enhance VirB binding to its site over that of the negative control ligand, UTP. In these assays, discrete shifts of the target DNA were not observed when VirB was added. Instead, VirB-DNA complexes appeared as smears (Fig. 1), suggesting that they were heterogeneous in nature. This was odd because the 54 bp DNA target is predicted to accommodate only two adjacent VirB dimers at most. Since high concentrations of BSA were included in these reactions to mitigate VirB-VirB interactions, the observed smears were unlikely to be VirB dimers bridging different DNA targets. Instead, the smears were more likely caused by VirB sliding off the linear DNA once it had specifically engaged its site. This speculation is supported by structural predictions, which show VirB forms a sliding clamp on DNA, where the DNA helix runs through the central lumen of the VirB dimer.

To further test these ideas, we next opted to use plasmid DNA in our EMSA experiments, a situation more relevant for VirB-DNA interactions *in vivo* and one that might allow us to capture VirB in its sliding clamp conformation on the DNA. Indeed, our experiments using closed circular plasmid DNA targets to study VirB-CTP-DNA interactions revealed that CTP allows VirB to form large molecular weight complexes on plasmid DNA, consistent with VirB sliding along the DNA after it has engaged its DNA recognition site. In contrast, the negative control, UTP, only allowed VirB to generate a small, discrete shift, likely caused by VirB engaging and remaining bound to its DNA recognition site. The finding that UTP generated a small, discrete shift in plasmid DNA (Fig. 2A), but not linear DNA (Fig. 1), was curious. It raised the possibility that VirB does not engage its site on linear relaxed DNA in the same way or with the same affinity as on supercoiled DNA. This is supported by VirB preferentially binding and shifting the supercoiled population of DNA in our plasmid EMSAs (the linear or relaxed plasmid molecules remain unchanged). Preliminary calculations reveal the affinity of VirB for its target site is around 7-fold higher on supercoiled plasmid DNA than on short 54 bp linear fragments (see Methods & relevant Figure legends). Based on these findings, and because VirB is unlikely to interact with linear DNA *in vivo*, we argue that there is limited use in studying VirB-DNA interactions on short, linear DNA fragments. Instead, we advocate for using supercoiled plasmid DNA in future studies aimed at elucidating how VirB-DNA interactions occur and how VirB functions *in vivo*.

In our plasmid EMSAs, regardless of whether CTP or UTP was present, other large discrete complexes were observed at the top of these gels. Initially, these were dismissed as an artifact of the plasmid EMSAs, which did not contain non-specific competitor DNA nor carrier protein, BSA. Importantly, though, after heat denaturation and subsequent DNA retrieval, we observed that these complexes were recovered differently (Fig. 2C). This suggests that the properties of the large discrete complexes formed in the presence of CTP and UTP differ. We speculate that the resilient high molecular weight complexes formed in the presence of CTP may be VirB-VirB bridged complexes. This is supported by ParB literature, where ParB-ParB bridges form in the presence of CTP (49), and by our work, which shows that GFP-VirB fusions form foci in the bacterial cytoplasm (50). Nevertheless, even if VirB is capable of forming bridged complexes, evidence supporting the involvement of VirB bridge formation in the mechanism of VirB-dependent transcriptional anti-silencing is lacking. Instead, transcriptional anti-silencing by VirB is demonstrated to rely on three key steps: i) the specific binding of dimeric VirB to its DNA recognition site (39); ii) the spreading of VirB dimers along the DNA (40); and iii) the VirB-mediated loss of negative supercoils in the region bound by H-NS (41).

Our previous work shows that VirB causes a transient loss of negative supercoiling in plasmid DNA when it binds to its DNA recognition site both *in vivo* and *in vitro*. *In vivo*, a localized loss of negative supercoiling, generated independently of VirB, can alleviate H-NS-mediated repression of a nearby promoter (41,44), implicating the VirB-dependent modulation of supercoiling in the mechanism of transcriptional anti-silencing. To study the role of CTP in the VirB-dependent modulation of DNA supercoiling, we exploited a set of VirB mutants with amino acid substitutions in residues predicted to contact CTP or in residues immediately adjacent to these. We found that VirB mutants unable to bind CTP (Fig. S3A), but able to bind the VirB DNA recognition site (Fig. S3B), do not trigger VirB-dependent modulation of DNA supercoiling (Fig. 3), providing the first evidence that CTP is required for the VirB-dependent loss of negative supercoiling, reported previously (41). Since CTP is necessary for the large complex formation and changes in DNA supercoiling, these two VirB-dependent activities likely fall in the same mechanistic pathway. Future work focused on probing the extent of VirB spread and how this impacts VirB-dependent changes in DNA topology on the large virulence plasmid would provide additional insight into the mechanism of anti-silencing in *Shigella*.

Taken together, our findings, along with the current understanding of other members of the ParB superfamily, allow us to present a new *in vivo* model of VirB-dependent transcriptional anti-silencing (Fig. 4). While VirB binds its DNA site in the absence of CTP (Fig. 1 and Fig. S1), *E. coli* (and likely *Shigella*) CTP pools are estimated to be ∼500 μM (51,52). Since the K_d_ values for VirB binding CTP are 9.3-14.3 μM (29,42,43), we posit that, *in vivo*, VirB binds CTP before engaging its site. Once bound to its recognition site, CTP is necessary for VirB to undergo a conformational change, where it adopts a DNA sliding clamp conformation (42,43). In this configuration, VirB spreads away from its binding site (40,43,48), allowing additional VirB dimers to dock to the now liberated site, leading to large VirB-DNA complexes to form on the DNA. These complexes cause a localized and transient loss of negative supercoils, which contributes to the remodeling of H-NS DNA complexes, thus leading to the VirB-mediated transcriptional anti-silencing of *Shigella* virulence genes (41).

**Figure 4.**
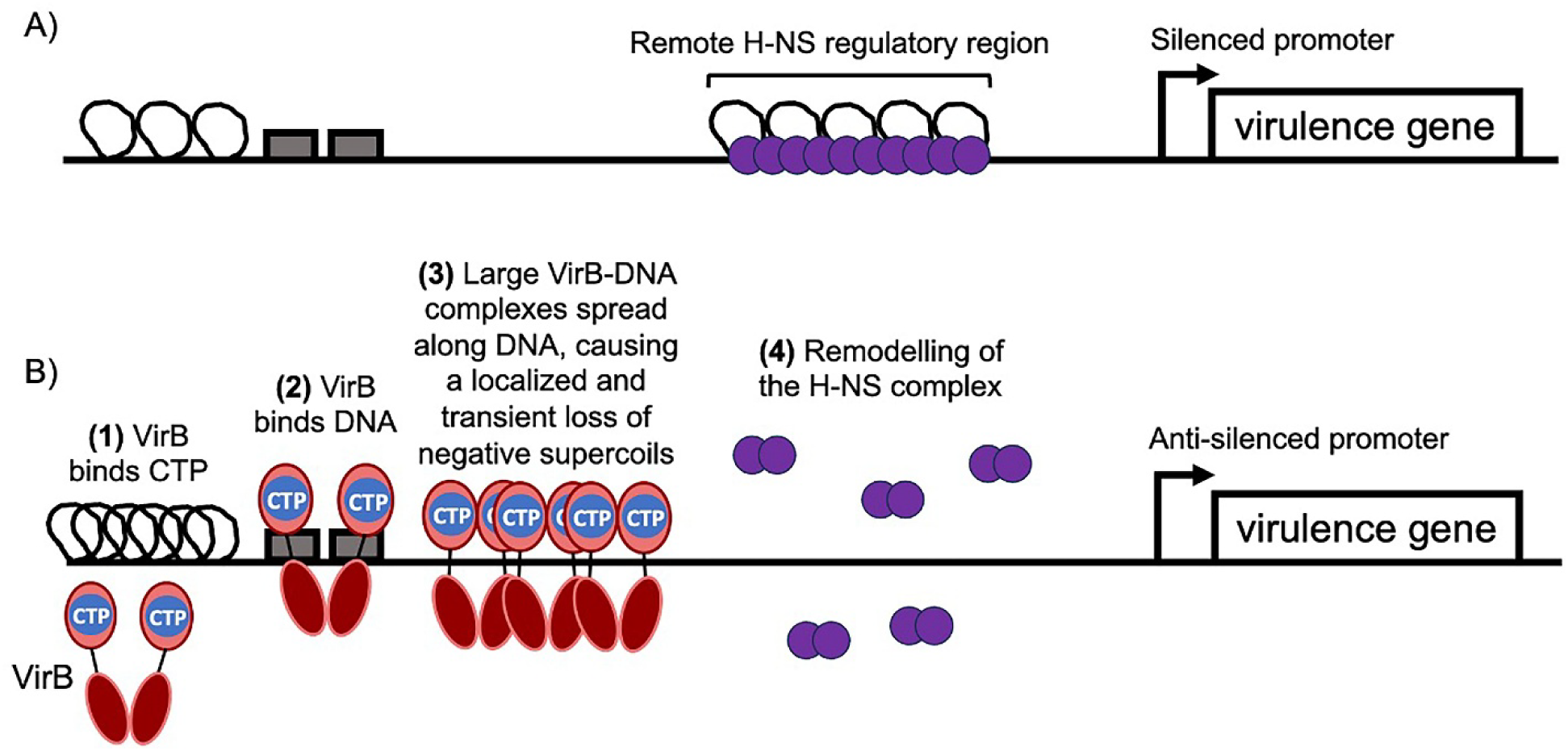
Model showing the mechanism of VirB-dependent anti-silencing *in vivo*. A) At 30°C, H-NS, often bound to regulatory regions upstream of the TSS, transcriptionally silences virulence genes (40,58). B) Entry into the host and a shift to 37°C leads to production of VirB. Once produced, VirB binds CTP at a 1:1 ratio (29), and dimeric VirB specifically engages the VirB binding site (9,39,40). Next, dimeric VirB undergoes a conformational change to form a DNA sliding clamp, allowing multiple VirB dimers to spread along DNA to form large VirB-DNA complexes. This results in the loss of negative supercoils in the region bound by H-NS, leading to the remodeling of the H-NS-DNA complex (41), which causes anti-silencing of virulence genes. **ALT TEXT:** Model of VirB-dependent anti-silencing *in vivo,* labelled A to B. A, shows H-NS silencing of virulence genes. B, shows the role that CTP plays in VirB-dependent anti-silencing and is presented as a numbered scheme (1) VirB binds CTP, (2) VirB binds DNA, (3) Large VirB-DNA complexes spread along DNA, causing a localized and transient loss of negative supercoils, (4) Remodelling of the H-NS complex.

Our proposed model of VirB-dependent transcriptional anti-silencing, where VirB slides along DNA in a sliding clamp conformation, is consistent with: i) structural predictions of VirB; ii) our finding that the large heat-resistant VirB DNA complexes form in plasmid EMSAs; and iii) DNA roadblocks can interfere with the anti-silencing activity by VirB (40). The ability of VirB to counteract H-NS-mediated silencing from remote sites (39,40,48) can also be reconciled by our model. One aspect of our model that is more challenging to explain is how VirB dimers locked onto DNA in a closed sliding-clamp conformation are released back to the cytoplasmic pool. In chromosomal ParB proteins, CTP hydrolysis reopens the DNA sliding clamp and removes ParB from the DNA (53–55). Recent work demonstrates that CTP hydrolysis by VirB is weaker than for chromosomal ParBs (42,43). So, currently, the mechanism of VirB release and the effect that low CTP hydrolysis rates have on the regulatory activities of VirB remain unclear. Work to address these two aspects of VirB-CTP-DNA interactions and their involvement in the VirB-dependent mechanism of anti-silencing is underway.

In sum, this work provides novel insight into VirB, a key regulator of *Shigella* virulence. The data presented in this study, along with previous research from our team (26,40,41) and others (25,42,43,56), support a mechanistic model for VirB-dependent transcriptional anti-silencing (Fig. 4) that now incorporates the role of its unusual ligand CTP. This work also raises additional questions about why CTP has evolved to modulate transcriptional anti-silencing in *Shigella* and whether fluctuations in the CTP pool size ever impact the activity of VirB. Additionally, it is interesting to consider if other plasmid-borne ParB-like proteins utilize CTP for different cellular functions. In this regard, recent work reveals several divergent bacterial, archaeal, and phage proteins may also bind CTP (57). Future studies investigating how CTP ligands are used by these and other unusual members of the ParB superfamily are clearly needed. These studies will further reveal the importance of CTP as an essential ligand for protein function.

## Data availability

The data supporting the findings of this study are available within the article, supplementary materials, or can be found in the following data repository: https://doi.org/10.5281/zenodo.17509235

## Supplementary Data Statement

Supplementary Data are available at *NAR* Online.

## Acknowledgements

We thank Monserate Biotechnology Group for the purification of VirB-His6. Special thanks to Elizabeth Huezo for her comments and edits, and to past and present lab members for insightful discussions.

## References

1. Dorman, C.J. (2009) Nucleoid-associated proteins and bacterial physiology. Adv Appl Microbiol, 67, 47–64.

2. Dorman, C.J. and Ni Bhriain, N. (1993) DNA topology and bacterial virulence gene regulation. Trends Microbiol, 1, 92–99.

3. Freddolino, P.L., Amemiya, H.M., Goss, T.J. and Tavazoie, S. (2022) Correction: Dynamic landscape of protein occupancy across the Escherichia coli chromosome. PLoS Biol, 20, e3001557.

4. Freddolino, L., Amemiya, H.M., Goss, T.J. and Tavazoie, S. (2021) Dynamic landscape of protein occupancy across the *Escherichia coli* chromosome. PLoS Biol, 19, e3001306.

5. Hengge-Aronis, R. (1999) Interplay of global regulators and cell physiology in the general stress response of *Escherichia coli*. Curr Opin Microbiol, 2, 148–152.

6. Falconi, M., Brandi, A., La Teana, A., Gualerzi, C.O. and Pon, C.L. (1996) Antagonistic involvement of FIS and H-NS proteins in the transcriptional control of hns expression. Mol Microbiol, 19, 965–975.

7. Falconi, M., Prosseda, G., Giangrossi, M., Beghetto, E. and Colonna, B. (2001) Involvement of FIS in the H-NS-mediated regulation of *virF* gene of *Shigella* and enteroinvasive *Escherichia coli*. Mol Microbiol, 42, 439–452.

8. O’Byrne, C.P. and Dorman, C.J. (1994) Transcription of the *Salmonella typhimurium spv* virulence locus is regulated negatively by the nucleoid-associated protein H-NS. FEMS Microbiol Lett, 121, 99–105.

9. Beloin, C., McKenna, S. and Dorman, C.J. (2002) Molecular dissection of VirB, a key regulator of the virulence cascade of *Shigella flexneri*. J Biol Chem, 277, 15333–15344.

10. Westermark, M., Oscarsson, J., Mizunoe, Y., Urbonaviciene, J. and Uhlin, B.E. (2000) Silencing and activation of ClyA cytotoxin expression in Escherichia coli. J Bacteriol, 182, 6347–6357.

11. Cathelyn, J.S., Ellison, D.W., Hinchliffe, S.J., Wren, B.W. and Miller, V.L. (2007) The RovA regulons of Yersinia enterocolitica and Yersinia pestis are distinct: evidence that many RovA-regulated genes were acquired more recently than the core genome. Mol Microbiol, 66, 189–205.

12. Yu, R.R. and DiRita, V.J. (2002) Regulation of gene expression in Vibrio cholerae by ToxT involves both antirepression and RNA polymerase stimulation. Mol Microbiol, 43, 119–134.

13. Yang, J., Hart, E., Tauschek, M., Price, G.D., Hartland, E.L., Strugnell, R.A. and Robins-Browne, R.M. (2008) Bicarbonate-mediated transcriptional activation of divergent operons by the virulence regulatory protein, RegA, from *Citrobacter rodentium*. Mol Microbiol, 68, 314–327.

14. Chen, C.C., Chou, M.Y., Huang, C.H., Majumder, A. and Wu, H.Y. (2005) A cis-spreading nucleoprotein filament is responsible for the gene silencing activity found in the promoter relay mechanism. J Biol Chem, 280, 5101–5112.

15. Chen, C.C. and Wu, H.Y. (2005) LeuO protein delimits the transcriptionally active and repressive domains on the bacterial chromosome. J Biol Chem, 280, 15111–15121.

16. Stoebel, D.M., Free, A. and Dorman, C.J. (2008) Anti-silencing: overcoming H-NS-mediated repression of transcription in Gram-negative enteric bacteria. Microbiology (Reading), 154, 2533–2545.

17. Hawkins, H.P. (1909) The identity of British ulcerative colitis and tropical bacillary dysentery. Br Med J, 2, 1331–1332.

18. Labrec, E.H., Schneider, H., Magnani, T.J. and Formal, S.B. (1964) Epithelial cell penetration as an essential step in the pathogenesis of bacillary dysentery. J Bacteriol, 88, 1503–1518.

19. Niyogi, S.K. (2007) Increasing antimicrobial resistance--an emerging problem in the treatment of shigellosis. Clin Microbiol Infect, 13, 1141–1143.

20. Niyogi, S.K. (2005) Shigellosis. J Microbiol, 43, 133–143.

21. Beloin, C. and Dorman, C.J. (2003) An extended role for the nucleoid structuring protein H-NS in the virulence gene regulatory cascade of *Shigella flexneri*. Mol Microbiol, 47, 825–838.

22. Gall, T.L., Mavris, M., Martino, M.C., Bernardini, M.L., Denamur, E. and Parsot, C. (2005) Analysis of virulence plasmid gene expression defines three classes of effectors in the type III secretion system of *Shigella flexneri*. Microbiology (Reading), 151, 951–962.

23. Porter, M.E. and Dorman, C.J. (1997) Differential regulation of the plasmid-encoded genes in the *Shigella flexneri* virulence regulon. Mol Gen Genet, 256, 93–103.

24. Basta, D.W., Pew, K.L., Immak, J.A., Park, H.S., Picker, M.A., Wigley, A.F., Hensley, C.T., Pearson, J.S., Hartland, E.L. and Wing, H.J. (2013) Characterization of the *ospZ* promoter in *Shigella flexneri* and its regulation by VirB and H-NS. J Bacteriol, 195, 2562–2572.

25. Turner, E.C. and Dorman, C.J. (2007) H-NS antagonism in *Shigella flexneri* by VirB, a virulence gene transcription regulator that is closely related to plasmid partition factors. J Bacteriol, 189, 3403–3413.

26. Wing, H.J., Yan, A.W., Goldman, S.R. and Goldberg, M.B. (2004) Regulation of IcsP, the outer membrane protease of the *Shigella* actin tail assembly protein IcsA, by virulence plasmid regulators VirF and VirB. J Bacteriol, 186, 699–705.

27. Tobe, T., Yoshikawa, M., Mizuno, T. and Sasakawa, C. (1993) Transcriptional control of the invasion regulatory gene *virB* of *Shigella flexneri*: activation by *virF* and repression by H-NS. J Bacteriol, 175, 6142–6149.

28. Taniya, T., Mitobe, J., Nakayama, S., Mingshan, Q., Okuda, K. and Watanabe, H. (2003) Determination of the InvE binding site required for expression of IpaB of the *Shigella sonnei* virulence plasmid: involvement of a ParB boxA-like sequence. J Bacteriol, 185, 5158–5165.

29. Gerson, T.M., Ott, A.M., Karney, M.M.A., Socea, J.N., Ginete, D.R., Iyer, L.M., Aravind, L., Gary, R.K. and Wing, H.J. (2023) VirB, a key transcriptional regulator of *Shigella* virulence, requires a CTP ligand for its regulatory activities. mBio, 14, e0151923.

30. Ireton, K. and Grossman, A.D. (1994) DNA-related conditions controlling the initiation of sporulation in *Bacillus subtilis*. Cell Mol Biol Res, 40, 193–198.

31. Mohl, D.A., Easter, J., Jr. and Gober, J.W. (2001) The chromosome partitioning protein, ParB, is required for cytokinesis in *Caulobacter crescentus*. Mol Microbiol, 42, 741–755.

32. McLean, T.C. and Le, T.B. (2023) CTP switches in ParABS-mediated bacterial chromosome segregation and beyond. Curr Opin Microbiol, 73, 102289.

33. Soh, Y.M., Davidson, I.F., Zamuner, S., Basquin, J., Bock, F.P., Taschner, M., Veening, J.W., De Los Rios, P., Peters, J.M. and Gruber, S. (2019) Self-organization of *parS* centromeres by the ParB CTP hydrolase. Science, 366, 1129-+.

34. Jalal, A.S.B., Tran, N.T., Wu, L.J., Ramakrishnan, K., Rejzek, M., Gobbato, G., Stevenson, C.E.M., Lawson, D.M., Errington, J. and Le, T.B.K. (2021) CTP regulates membrane-binding activity of the nucleoid occlusion protein Noc. Mol Cell, 81, 3623–3636 e3626.

35. Osorio-Valeriano, M., Altegoer, F., Steinchen, W., Urban, S., Liu, Y., Bange, G. and Thanbichler, M. (2019) ParB-type DNA Segregation Proteins Are CTP-Dependent Molecular Switches. Cell, 179, 1512-+.

36. Jalal, A.S., Tran, N.T. and Le, T.B. (2020) ParB spreading on DNA requires cytidine triphosphate in vitro. Elife, 9.

37. Babl, L., Giacomelli, G., Ramm, B., Gelmroth, A.K., Bramkamp, M. and Schwille, P. (2022) CTP-controlled liquid-liquid phase separation of ParB. J Mol Biol, 434, 167401.

38. Szymczak, J., Strzalka, A., Bania, D., Jakimowicz, D. and Szafran, M.J. (2025) Significance of the CTP-binding motif for the interactions of S. coelicolor ParB with DNA, chromosome segregation, and sporogenic hyphal growth. Nucleic Acids Res, 53.

39. Castellanos, M.I., Harrison, D.J., Smith, J.M., Labahn, S.K., Levy, K.M. and Wing, H.J. (2009) VirB alleviates H-NS repression of the *icsP* promoter in *Shigella flexneri* from sites more than one kilobase upstream of the transcription start site. J Bacteriol, 191, 4047–4050.

40. Weatherspoon-Griffin, N., Picker, M.A., Pew, K.L., Park, H.S., Ginete, D.R., Karney, M.M., Usufzy, P., Castellanos, M.I., Duhart, J.C., Harrison, D.J. et al. (2018) Insights into transcriptional silencing and anti-silencing in *Shigella flexneri:* a detailed molecular analysis of the *icsP* virulence locus. Mol Microbiol, 108, 505–518.

41. Picker, M.A., Karney, M.M.A., Gerson, T.M., Karabachev, A.D., Duhart, J.C., McKenna, J.A. and Wing, H.J. (2023) Localized modulation of DNA supercoiling, triggered by the *Shigella* anti-silencer VirB, is sufficient to relieve H-NS-mediated silencing. Nucleic Acids Res.

42. Antar, H. and Gruber, S. (2023) VirB, a transcriptional activator of virulence in *Shigella flexneri*, uses CTP as a cofactor. Commun Biol, 6, 1204.

43. Jakob, S., Steinchen, W., Hanssmann, J., Rosum, J., Langenfeld, K., Osorio-Valeriano, M., Steube, N., Giammarinaro, P.I., Hochberg, G.K.A., Glatter, T. et al. (2024) The virulence regulator VirB from *Shigella flexneri* uses a CTP-dependent switch mechanism to activate gene expression. Nat Commun, 15, 318.

44. Karney, M.M.A., Gerson, T.M., Picker, M.A. and Wing, H.J. (2024) Methods to quantitatively measure topological changes induced by DNA-binding proteins in vivo and in vitro. Methods Mol Biol, 2819, 421–441.

45. Guzman, L.M., Belin, D., Carson, M.J. and Beckwith, J. (1995) Tight regulation, modulation, and high-level expression by vectors containing the arabinose PBAD promoter. J Bacteriol, 177, 4121–4130.

46. Gao, X., Zou, T., Mu, Z., Qin, B., Yang, J., Waltersperger, S., Wang, M., Cui, S. and Jin, Q. (2013) Structural insights into VirB-DNA complexes reveal mechanism of transcriptional activation of virulence genes. Nucleic Acids Res, 41, 10529–10541.

47. Venkatesan, M.M., Goldberg, M.B., Rose, D.J., Grotbeck, E.J., Burland, V. and Blattner, F.R. (2001) Complete DNA sequence and analysis of the large virulence plasmid of *Shigella flexneri*. Infect Immun, 69, 3271–3285.

48. Cris, C., Karney, M.M.A., Rosen, J.S., Karabachev, A.D., Huezo, E.N. and Wing, H.J. (2025) Remote regulation by VirB, the transcriptional anti-silencer of *Shigella* virulence genes, provides mechanistic information. Mol Microbiol, 123, 265–278.

49. Tisma, M., Panoukidou, M., Antar, H., Soh, Y.M., Barth, R., Pradhan, B., Barth, A., van der Torre, J., Michieletto, D., Gruber, S., et al. (2022) ParB proteins can bypass DNA-bound roadblocks via dimer-dimer recruitment. Sci Adv, 8, eabn3299.

50. Socea, J.N., Bowman, G.R. and Wing, H.J. (2021) VirB, a key transcriptional regulator of virulence plasmid genes in *Shigella flexneri*, forms DNA-binding site-dependent foci in the bacterial cytoplasm. J Bacteriol, 203.

51. Buckstein, M.H., He, J. and Rubin, H. (2008) Characterization of nucleotide pools as a function of physiological state in *Escherichia coli*. J Bacteriol, 190, 718–726.

52. Zbornikova, E., Knejzlik, Z., Hauryliuk, V., Krasny, L. and Rejman, D. (2019) Analysis of nucleotide pools in bacteria using HPLC-MS in HILIC mode. Talanta, 205, 120161.

53. Antar, H., Soh, Y.M., Zamuner, S., Bock, F.P., Anchimiuk, A., De los Rios, P. and Gruber, S. (2021) Relief of ParB autoinhibition by *parS* DNA catalysis and recycling of ParB by CTP hydrolysis promote bacterial centromere assembly. Sci Adv, 7.

54. Osorio-Valeriano, M., Altegoer, F., Das, C.K., Steinchen, W., Panis, G., Connolley, L., Giacomelli, G., Feddersen, H., Corrales-Guerrero, L., Giammarinaro, P.I. et al. (2021) The CTPase activity of ParB determines the size and dynamics of prokaryotic DNA partition complexes. Mol Cell, 81, 3992–4007 e3910.

55. Jalal, A.S., Tran, N.T., Stevenson, C.E., Chimthanawala, A., Badrinarayanan, A., Lawson, D.M. and Le, T.B. (2021) A CTP-dependent gating mechanism enables ParB spreading on DNA. Elife, 10.

56. McKenna, S., Beloin, C. and Dorman, C.J. (2003) *In vitro* DNA-binding properties of VirB, the *Shigella flexneri* virulence regulatory protein. FEBS Lett, 545, 183–187.

57. Kaljevic, J., Sukhoverkov, K.V., Johnson, K.E., Hocher, A. and Le, T.B.K. (2025) Versatile NTP recognition and domain fusions expand the functional repertoire of the ParB-CTPase fold beyond chromosome segregation. Proc Natl Acad Sci U S A, 122, e2527592122.

58. McKenna, J.A. and Wing, H.J. (2020) The Antiactivator of Type III Secretion, OspD1, Is Transcriptionally Regulated by VirB and H-NS from Remote Sequences in *Shigella flexneri*. J Bacteriol, 202.

