## Supplementary Materials for "The role of the VirB ligand CTP in the molecular mechanism of transcriptional anti-silencing in *Shigella flexneri*"

### Supplemental Materials and Methods

#### Differential radial capillary action of ligand assay

Differential radial capillary action of ligand assay, DRaCALA, was adapted from references (1-4) and performed as described in (5). Briefly, radiolabeled (hot) $\alpha^{32}$P-CTP (5 nM final) was prepared in buffer containing 500 mM NaCl, 500 mM Tris/HCl, pH 7.5, 50 mM $MgCl_2$, and kept on ice. Purified VirB protein (25 µM final) was added to each reaction and incubated for 10 minutes at room temperature. After incubation, 4 µL were spotted onto dry nitrocellulose (GE Healthcare) in three technical replicates, allowed to air dry, and then exposed to a storage phosphor screen (Kodak). The screen was imaged on an Amersham Typhoon. Ligand binding was indicated by a darker inner core staining, due to rapid protein immobilization. Spots were quantified by densitometric analysis of the inner and outer core of each spot using AzureSpot Analysis Software version 2.0.062. The “Analysis Toolbox” was used to draw circles around the outer and inner cores of each spot. Circles were copied using the “duplicate” feature and dragged to adjacent spots to allow for direct comparisons. The densitometric output for each spot was used to calculate the fraction bound ($F_B$) using the following equation: $F_B = (I_{inner} – (A_{inner} \times [(I_{total} - I_{inner}) / (A_{total} - A_{inner})]))/I_{total}$ (1).

**Table S1. Bacterial strains and plasmids used in this study**

| Label | Description | Reference |
|---|---|---|
| **Strains** | | |
| *E. coli* | | |
| DH10B | F- endA1 deoR+ recA1 galE15 galK16 nupG rpsL Δ(lac)X74 φ80lacZΔM15 araD139 Δ(ara, leu)7697 mcrA Δ(mrr-hsdRMS-mcrBC) StrR λ– | (6) |
| M15 (pREP4) | *E. coli* strain K-12; F-, Φ80ΔlacM15, thi, lac-, mtl-, recA+, $Km^r$ | Qiagen |
| **Plasmids** | | |
| pATM324 | pBAD18-*virB*; $Amp^r$ | (7) |
| pQE60 | Cloning vector for production of C-terminally His–tagged proteins | Qiagen |
| pBAD18 | Arabinose-inducible pBAD expression vector, ori pBR; $Amp^r$ | (8) |
| pTMG24 | pBAD-*virB* K152E-R167E; $Amp^r$ | (5) |
| pDRG04 | pBAD-*virB* R94A; $Amp^r$ | (5) |
| pDRG05 | pBAD-*virB* R95A; Ampr | (5) |
| pAMO12 | pBAD-*virB* T68A; $Amp^r$ | (5) |
| pAMO13 | pBAD-*virB* T68S; $Amp^r$ | (5) |
| pAJH01 | pQE60-*virB*-His6-tagged; $Amp^r$ | This work |

**Table S2. Complete statistics for linear EMSA with WT and MUT VirB binding site**

| Linear EMSA – WT site and CTP | | VirB concentration | | |
|---|---|---|---|---|
| | | 0 µM | 2 µM | 4 µM |
| VirB concentration | 0 µM | | <0.001* | <0.001* |
| | 2 µM | | | 0.106 |
| | 4 µM | | | |

| Linear EMSA – WT site and UTP | | VirB concentration | | |
|---|---|---|---|---|
| | | 0 µM | 2 µM | 4 µM |
| VirB concentration | 0 µM | | 0.795 | 0.042* |
| | 2 µM | | | 0.209 |
| | 4 µM | | | |

| Linear EMSA – MUT site and CTP | | VirB concentration | | |
|---|---|---|---|---|
| | | 0 µM | 2 µM | 4 µM |
| VirB concentration | 0 µM | | 1.000 | 1.000 |
| | 2 µM | | | 1.000 |
| | 4 µM | | | |

| Linear EMSA – MUT site and UTP | | VirB concentration | | |
|---|---|---|---|---|
| | | 0 µM | 2 µM | 4 µM |
| VirB concentration | 0 µM | | 1.000 | 0.528 |
| | 2 µM | | | 1.000 |
| | 4 µM | | | |

Significance was determined using a one-way ANOVA with post hoc Bonferroni. Asterisks indicate $p<0.05$. Grey boxes represent data that was not compared.

**Table S3. Complete Statistics for linear EMSA with varying VirB concentration**

| **Linear EMSA – Varying Protein Concentration (CTP condition)** | | **VirB concentration** | | | | | | |
|---|---|---|---|---|---|---|---|---|
| | | 0 µM | 0.25 µM | 0.5 µM | 0.675 µM | 0.75 µM | 1 µM | 4 µM |
| **VirB concentration** | 0 µM | | 0.987 | 0.85 | 0.499 | 0.066 | 0.001* | <0.001* |
| | 0.25 µM | | | 0.998 | 0.893 | 0.228 | 0.005* | <0.001* |
| | 0.5 µM | | | | 0.994 | 0.469 | 0.013* | <0.001* |
| | 0.675 µM | | | | | 0.826 | 0.041* | <0.001* |
| | 0.75 µM | | | | | | 0.359 | <0.001* |
| | 1 µM | | | | | | | 0.005* |
| | 4 µM | | | | | | | |

| **Linear EMSA – Varying Protein Concentration (UTP condition)** | | **VirB concentration** | | | | | | |
|---|---|---|---|---|---|---|---|---|
| | | 0 µM | 0.25 µM | 0.5 µM | 0.675 µM | 0.75 µM | 1 µM | 4 µM |
| **VirB concentration** | 0 µM | | 1 | 0.99 | 0.991 | 0.947 | 0.678 | 0.186 |
| | 0.25 µM | | | 1 | 1 | 0.995 | 0.869 | 0.324 |
| | 0.5 µM | | | | 1 | 0.998 | 0.908 | 0.372 |
| | 0.675 µM | | | | | 1 | 0.962 | 0.48 |
| | 0.75 µM | | | | | | 0.995 | 0.678 |
| | 1 µM | | | | | | | 0.938 |
| | 4 µM | | | | | | | |

Significance was determined using a one-way ANOVA with post hoc Tukey HSD. Asterisks indicate p<0.05. Grey boxes represent data that was not compared.

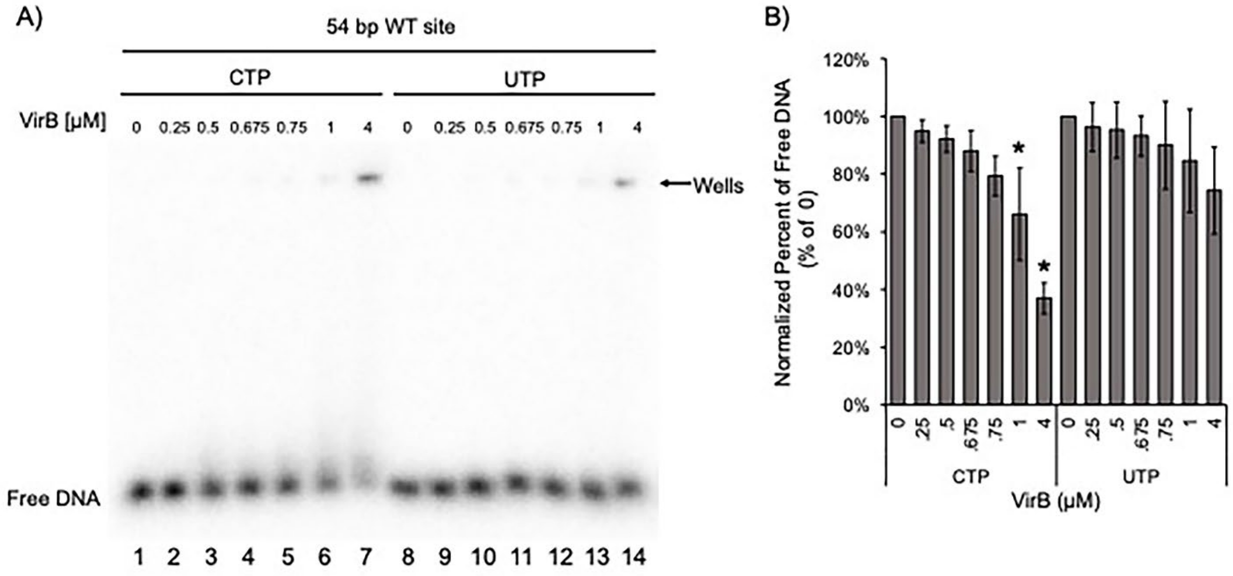

**Figure S1. Examining VirB interactions with a linear DNA using a finer titration scheme of VirB.** A) EMSA where target DNA containing a WT VirB binding site was incubated with increasing concentrations of VirB (between 0-1 and 4 μM) incubated in a molar excess of either CTP or UTP. B) Quantification of signal loss of the lower band in Fig. S1A, as VirB concentration increases. The molar ratio of VirB:DNA required to shift linear DNA was calculated to be 206:1. Significance was calculated using a one-way ANOVA post-hoc Tukey HSD, $p<0.05$. Asterisks, *, indicate a significant decrease compared to 0 lanes in each given condition. Complete statistical analysis provided in Table S3.

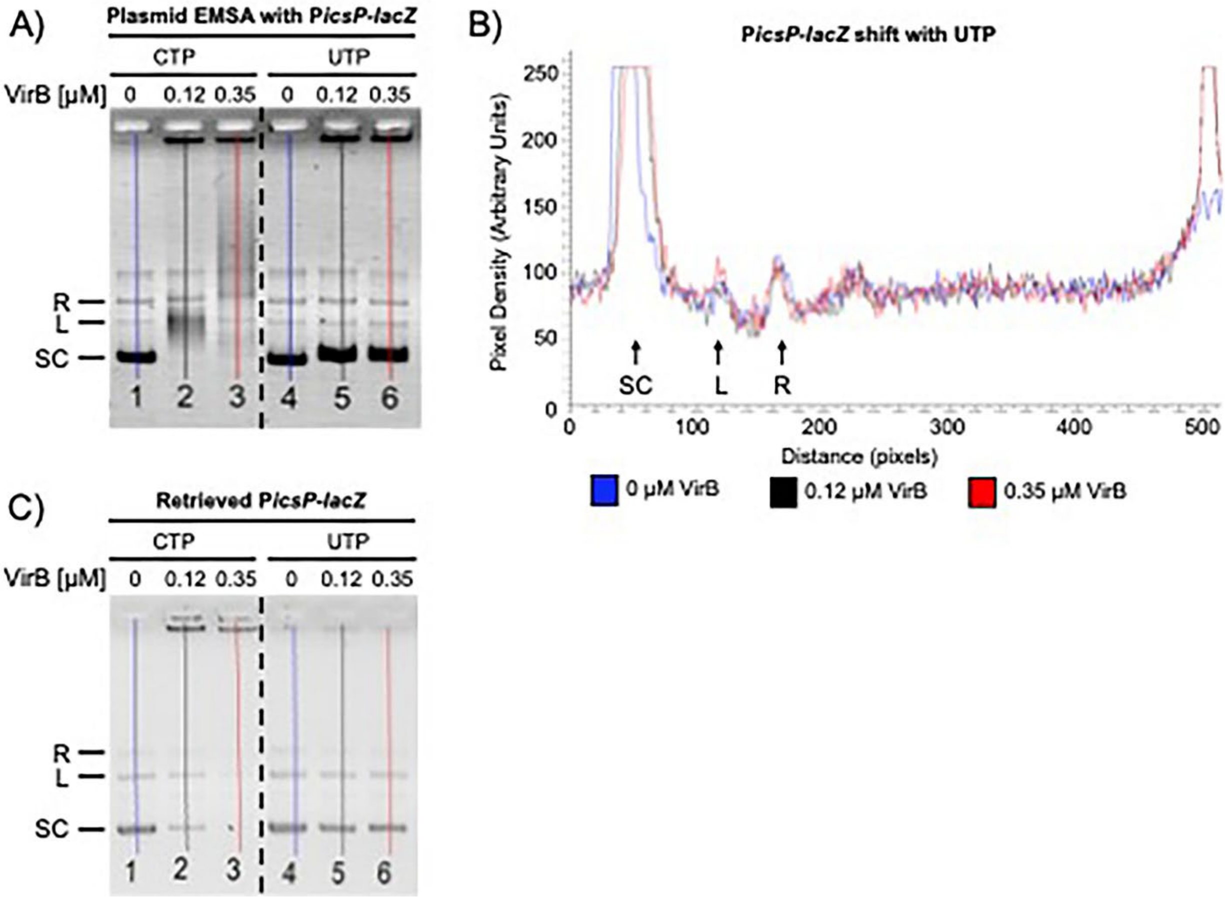

**Figure S2. Quantification of plasmid EMSAs and DNA capture assay.** A) Lane trace of plasmid EMSA with P*icsP-lacZ* (Fig. 2A) used to generate lane profiles in Fig. 2B and Fig. S2B. B) The lane profile of the shift of P*icsP-lacZ* in the presence of UTP (lanes 4-6) from Fig. 2A. C) Lane trace of DNA capture assay (Fig. 2C) used to generate lane profiles in Fig. 2D. The peaks corresponding to the supercoiled (SC), linear (L), and relaxed (R) bands are indicated with arrows.

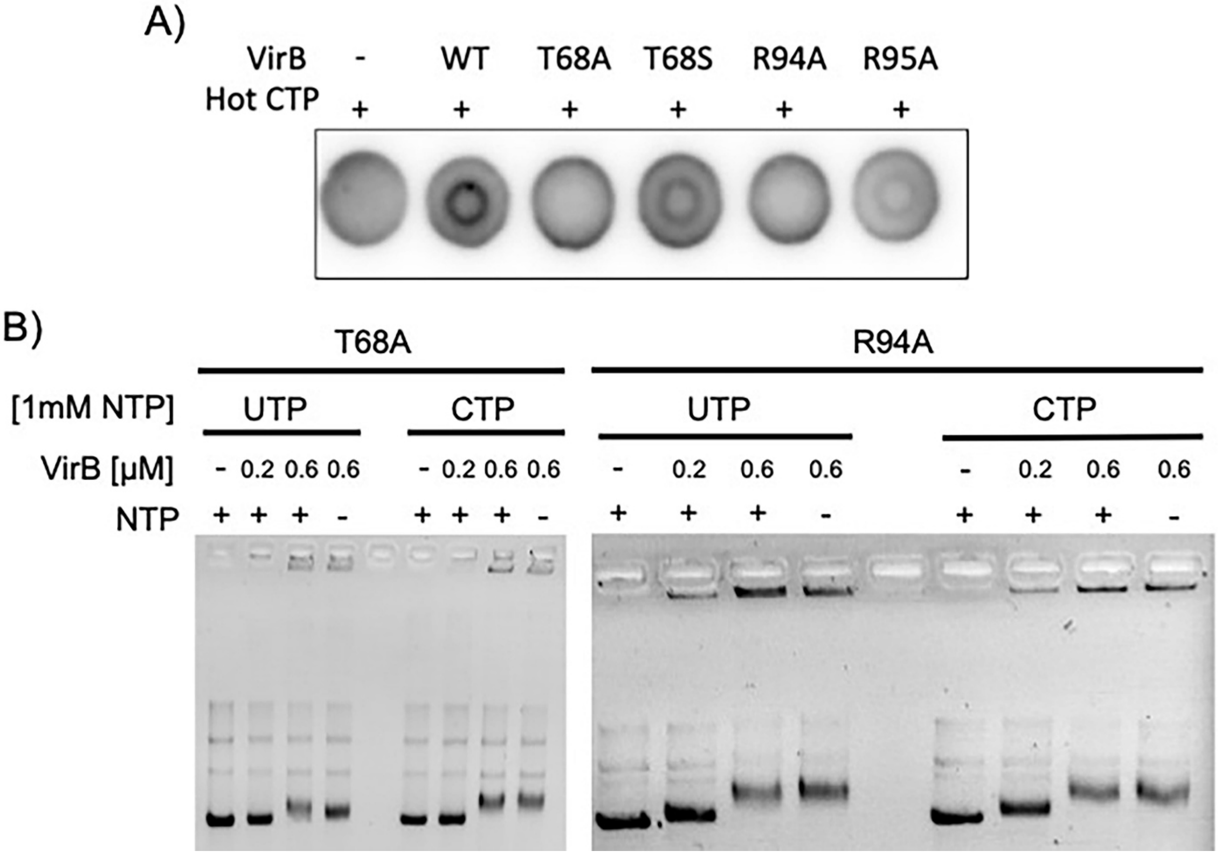

**Figure S3. Validation of VirB-CTP binding mutants.** A) DRaCALA images assessing the ability of 25 μM of VirB-His6 derivatives to bind to 5 nM α$^{32}$P-CTP. B) Modified electromobility shift assay with plasmid DNA containing a VirB binding site (P*icsP-lacZ)* with excess CTP or UTP (1mM) and increasing concentrations (0, 0.2, 0.6 μM) of VirB-His6 derivatives (T68A, or R94A).

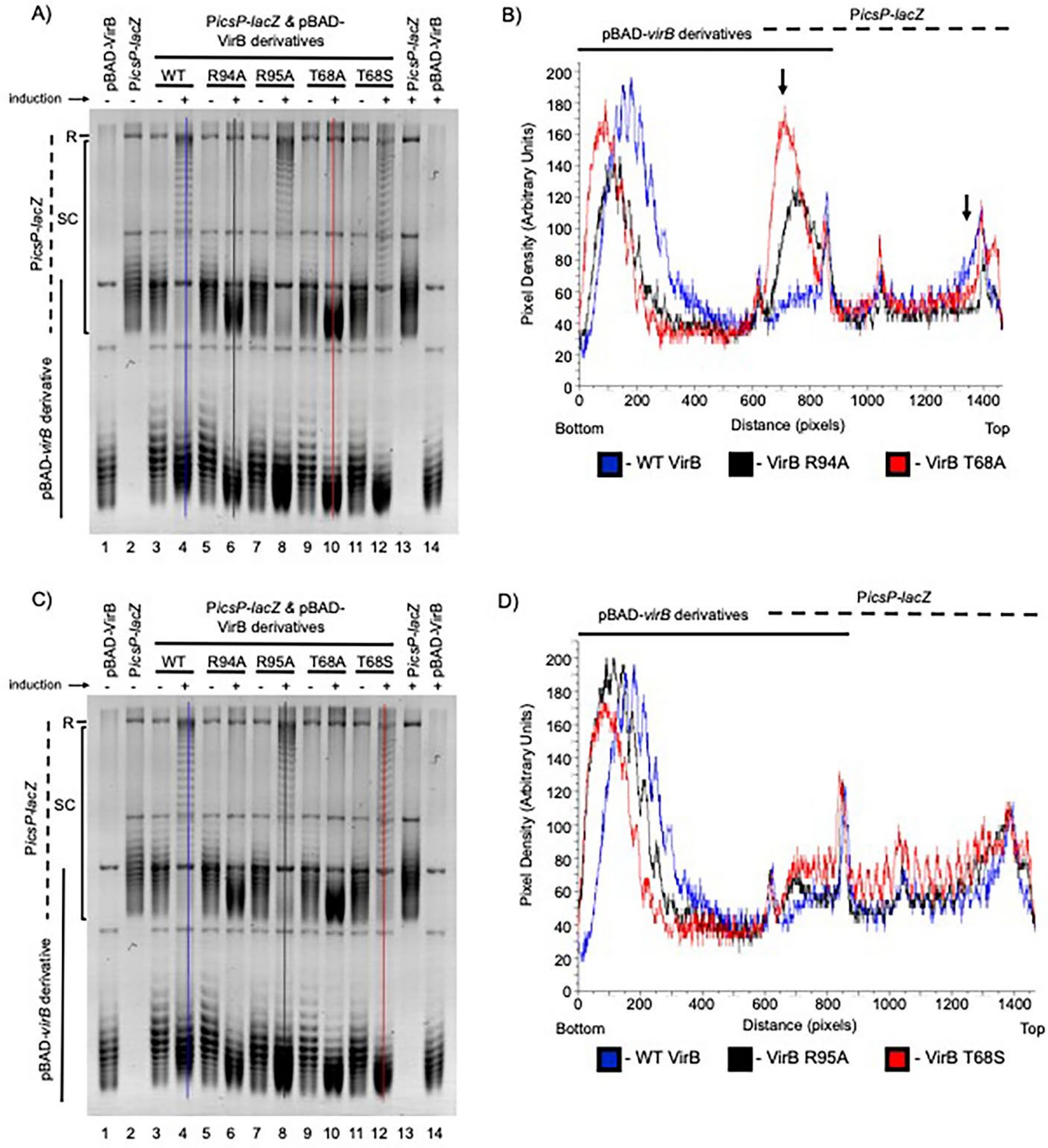

**Figure S4. Densitometry of *in vivo* VirB-dependent changes in DNA supercoiling using CTP binding mutants.** A) Lane trace analyses of WT, VirB R94A, and VirB T68A. B) Profile of lanes traces for WT, VirB R94A, and VirB T68A. C) Lane trace analyses of WT, VirB R95A, and VirB T68S. D) Profile of lanes traces for WT, VirB R95A, and VirB T68S. Lane traces were routinely drawn from the bottom to the top of the gel. In each case, lane trace analyses of three lanes with induction of VirB or VirB derivatives are shown to facilitate comparison of lanes of interest without signal crowding. Profiles of lane traces routinely display dashed lines to denote the signal corresponding to pBAD-*virB* or derivatives, and solid lines to denote the signal corresponding to P*icsP-lacZ.* For ease of interpretation, significant changes between topoisomer distributions in P*icsP-lacZ* in the presence of each VirB derivative are highlighted with downward-facing arrows in panel B.

### References

1. Roelofs, K.G., Wang, J., Sintim, H.O. and Lee, V.T. (2011) Differential radial capillary action of ligand assay for high-throughput detection of protein-metabolite interactions. *Proc Natl Acad Sci U S A*, **108**, 15528-15533.
2. Soh, Y.M., Davidson, I.F., Zamuner, S., Basquin, J., Bock, F.P., Taschner, M., Veening, J.W., De Los Rios, P., Peters, J.M. and Gruber, S. (2019) Self-organization of *parS* centromeres by the ParB CTP hydrolase. *Science*, **366**, 1129-+.
3. Osorio-Valeriano, M., Altegoer, F., Steinchen, W., Urban, S., Liu, Y., Bange, G. and Thanbichler, M. (2019) ParB-type DNA Segregation Proteins Are CTP-Dependent Molecular Switches. *Cell*, **179**, 1512-+.
4. Jalal, A.S., Tran, N.T. and Le, T.B. (2020) ParB spreading on DNA requires cytidine triphosphate in vitro. *Elife*, **9**.
5. Gerson, T.M., Ott, A.M., Karney, M.M.A., Socea, J.N., Ginete, D.R., Iyer, L.M., Aravind, L., Gary, R.K. and Wing, H.J. (2023) VirB, a key transcriptional regulator of *Shigella* virulence, requires a CTP ligand for its regulatory activities. *mBio*, **14**, e0151923.
6. Grant, S.G., Jessee, J., Bloom, F.R. and Hanahan, D. (1990) Differential plasmid rescue from transgenic mouse DNAs into *Escherichia coli* methylation-restriction mutants. *Proc Natl Acad Sci U S A*, **87**, 4645-4649.
7. Schuch, R. and Maurelli, A.T. (1997) Virulence plasmid instability in *Shigella flexneri* 2a is induced by virulence gene expression. *Infect Immun*, **65**, 3686-3692.
8. Guzman, L.M., Belin, D., Carson, M.J. and Beckwith, J. (1995) Tight regulation, modulation, and high-level expression by vectors containing the arabinose PBAD promoter. *J Bacteriol*, **177**, 4121-4130.
